# Cryo-EM structures reveal transcription initiation steps by yeast mitochondrial RNA polymerase

**DOI:** 10.1101/2020.04.13.038620

**Authors:** Brent De Wijngaert, Shemaila Sultana, Chhaya Dharia, Hans Vanbuel, Jiayu Shen, Daniel Vasilchuk, Sergio E. Martinez, Eaazhisai Kandiah, Smita S. Patel, Kalyan Das

## Abstract

Cryo-EM structures of transcription pre-initiation complex (PIC) and initiation complex (IC) of yeast mitochondrial RNA polymerase show fully resolved transcription bubbles and explain promoter melting, template alignment, DNA scrunching, transition into elongation, and abortive synthesis. Promoter melting initiates in PIC with MTF1 trapping the −4 to −2 non-template (NT) bases in its NT-groove. Transition to IC is marked by a large-scale movement that aligns the template with RNA at the active site. RNA synthesis scrunches the NT strand into an NT-loop, which interacts with centrally positioned MTF1 C-tail. Steric clashes of the C-tail with RNA:DNA and NT-loop, and dynamic scrunching-unscrunching of DNA explain abortive synthesis and transition into elongation. Capturing the catalytically active IC-state with UTPαS poised for incorporation enables modeling toxicity of antiviral nucleosides/nucleotides.

---

Gene transcription is catalyzed by DNA-dependent RNA polymerases (RNAPs) in a multi-step regulated process (*1–4*). Multi-subunit and single-subunit RNAPs catalyze transcription initiation by a series of events – (i) promoter-specific recognition, (ii) formation of a transcription bubble, (iii) RNA synthesis accompanied by DNA scrunching, and (iv) aborted synthesis of short RNA transcripts or transition to the elongation phase with promoter release. Understanding the mechanism of transcription initiation requires capturing key intermediate states of RNAP complexes and characterizing them biochemically and structurally.

Mitochondrial DNA is transcribed by single-subunit RNAPs (mtRNAP), which unlike their phage counterparts, depend on one or more transcription factors for promoter-specific transcription initiation. Much of our understanding of mitochondrial DNA transcription comes from studies of yeast (*S. cerevisiae*) and human mtRNAPs (*2, 3, 5*). The yeast mtRNAP transcription initiation complex (y-mtIC) is comprised of the catalytic subunit RPO41 and a transcription factor MTF1. The human mtRNAP transcription initiation complex (h-mtIC) comprises of POLRMT and two transcription factors. The h-mtIC has been structurally characterized by crystallography (*6*), but the structure was captured in an inactive fingers-clenched state with a major part of the transcription bubble disordered. Hence, the structural basis for promoter melting, DNA scrunching, and transcription initiation remains largely unknown for the mtRNAPs.

RPO41 (ΔN100) and MTF1 were assembled on a pre-melted promoter (−21 to +12, *15S* yeast mtDNA promoter; Fig. 1A) to generate the yeast mitochondrial transcription pre-initiation complex (y-mtPIC) (Fig. S1). The y-mtPIC was incubated with pppGpG RNA and a non-hydrolysable UTPαS to generate the y-mtIC poised to incorporate the +3 NTP. Single-particle cryo-EM data analysis of the quinary y-mtIC revealed a surprising coexistence of PIC and IC states in equilibrium (Fig. S2). The y-mtIC structure had bound RNA and NTP, and y-mtPIC structure had no RNA or NTP. Another dataset collected from a PIC-only grid extends the resolution to 3.1 Å. Key steps guiding transcription initiation are revealed from the 3.1 Å y-mtPIC and 3.7 Å y-mtIC structures.

**Fig. 1.**
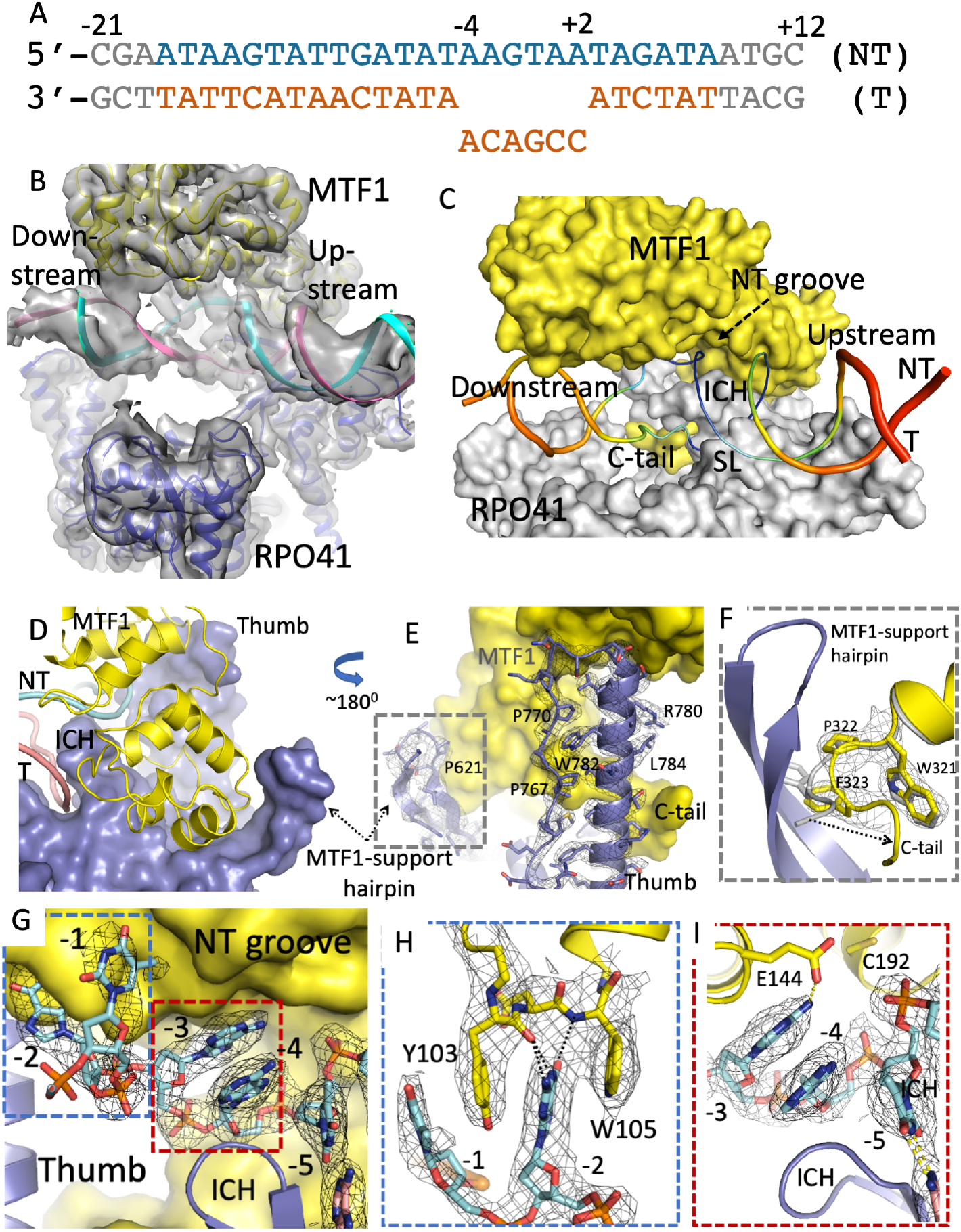
Structure of the yeast mitochondrial RNA polymerase transcription pre-initiation complex (y-mtPIC). **A.** Nucleotide sequence of the *15S* yeast mtDNA promoter with mutated template strand generates a pre-melted initiation bubble; missing nucleotides in gray. **B.** Unsharpened 3.1 Å resolution density map shows the DNA track in the y-mtPIC structure; blue RPO41, yellow MTF1, cyan non-template (NT), and pink template (T). **C**. B-factor putty (thin blue, lowest to thick red, highest B-factors) representation of DNA in the active-site cavity; MTF1 (yellow) and RPO41 (gray), intercalating hairpin (ICH), specificity loop (SL), non-template (NT) groove and Mtf1 C-tail are highlighted; see Fig. S3 for details. **D.** RPO41 thumb, ICH, and MTF1-supporting hairpin (613-632) place MTF1 (yellow ribbon) in y-mtPIC. **E.** The 3.1 Å resolution density map defines thumb-domain interactions with MTF1 (yellow surface). **F.** A zoomed view of the highlighted region in panel E shows the MTF1-supporting loop guiding the C-tail inward; the arrow shows the repositioning of C-tail base in Mft1-only structure (PDB Id. 1I4W; gray) to that in y-mtIC. **G.** The interactions of −4 to −1 NT nucleotides with the NT-groove in Mtf1 (yellow surface). The density map defines the track and interactions of the NT nucleotides; the boxed regions are elaborated in panels H and I. **H.** The −2 NT base has aromatic ring stacking and extensive base-specific interactions (stereo view, Fig S3B); N1 and N2 interact with the main-chain carbonyl of Y103, O6 with the main-chain amide of W105, and N3 and N9 interact with Oε1 of E145 sidechain. **I.** Position and interactions of −3 and −4 NT bases.

In y-mtPIC structure (Fig. 1B & C), the MTF1 is traced from 2 to 336 out of 341 amino acid residues, RPO41 is traced from 386 to the end residue 1351 with few disordered regions, and unambiguously traced DNA (Fig. S3A). A stable transcription core is composed of RPO41, MTF1, and transcription bubble (Fig. 1C). RPO41 interacts with MTF1 at multiple locations. Two RPO41 β-hairpins – the intercalating hairpin (ICH) and the MTF1-supporting hairpin (K613-P632) form a crescent-shaped platform that accommodates the C-terminal domain of MTF1 (252-325) (Fig. 1D). The N-terminal domain of MTF1 (2-251) contacts the tip of the RPO41 thumb helix (Fig. 1E); biochemically, we show the interaction stabilizes RPO41-MTF1 complex (Fig. S3C-D)(*7*). The MTF1-supporting hairpin also guides the MTF C-tail (326-341) towards the active site (Fig. 1F); the C-tail is disordered in free MTF1(*8*).

The PIC structure has not been observed previously; thus, it provides new insights into the mechanism of promoter melting. The y-mtPIC structure suggests that RPO41 and MTF1 initiate the promoter melting by creating a 4-nt transcription bubble from −4 to −1. We provided a −4 to +2 pre-melted promoter; however, the +1 and +2 nucleotides assume a duplex-like DNA conformation in PIC albeit lacking canonical base-pairing. DNA melting is driven by sequence-specific interactions of the NT strand with the NT-groove that lies at the interface of N- and C-terminal domains of MTF1 (residues 103-105, 144-148, and 190-192). The −4 to −2 AAG bases in the NT strand are flipped towards NT-groove (Fig. 1G). The −2 guanine base is sandwiched between the aromatic side chains of Y103 and W105, and all N and O atoms of the base, except N7, are engaged in complementary hydrogen bond interactions (Fig. 1H); mutation of −2 guanine severely impair promoter melting (*9*). The −1 NT base stacks with Y103 with no base-specific interaction. The −3 and −4 AA bases are stacked together and sandwiched by ICH on one side and an MTF1 helix (144-156) on the other side (Fig. 1I). Mutations of MTF1 residues near the promoter DNA, E144F, R178A+K179A (Fig. S3E-F), and C192F (*10*) lower the efficiency of transcription initiation. The structure reveals that the aromatic sidechain of mutated E144F or C192F interferes with the binding of NT −3 and −4 bases and R178A+K179A would reduce DNA-backbone interactions.

The quinary y-mtIC has a promoter designed to bind a 2-mer RNA and incorporate a third nucleotide (Fig. 2A). The 3.7 Å density map of IC locates many previously uncharacterized structural elements, including the C-tail of MTF1 and the scrunched DNA (Fig. 2B-C), and reveals the mechanism of template alignment and RNA synthesis during initiation. Comparison of PIC and IC shows little shift of upstream promoter DNA and interacting structural elements (including ICH, specificity loop, and thumb of RPO41 and NT-groove of MTF1) during PIC to IC transition. In contrast, the downstream DNA and interacting C-terminal domain of RPO41 undergo large conformational changes including fingers closing (Movie S1). During the transition, the upstream DNA from position −1 onward is locked in the MTF1 NT-groove while the template strand undergoes a large conformational switching to align with the 2-mer RNA and the incoming UTPαS at the active site (Fig. 2C; Movie S2). These unsynchronized events at two ends of the transcription bubble scrunches the NT strand into an NT-loop (Movie S3). DNA scrunching has been proposed in multi- and single-subunit RNAPs including y-mtRNAP (*11–14*). Our y-mtIC structure is first to capture the scrunched conformation (Fig. 2C). The scrunched NT-loop is stabilized by ICH (H641, N642), thumb (R780 and K787), and MTF1 C-tail (M334-Y335). The looping of NT strand alters the downstream DNA track and bends it from ~60° to ~120° with respect to the upstream DNA; thus, transforming a V-shaped DNA in PIC to an U-shape in IC (Fig. S5).

**Figure 2.**
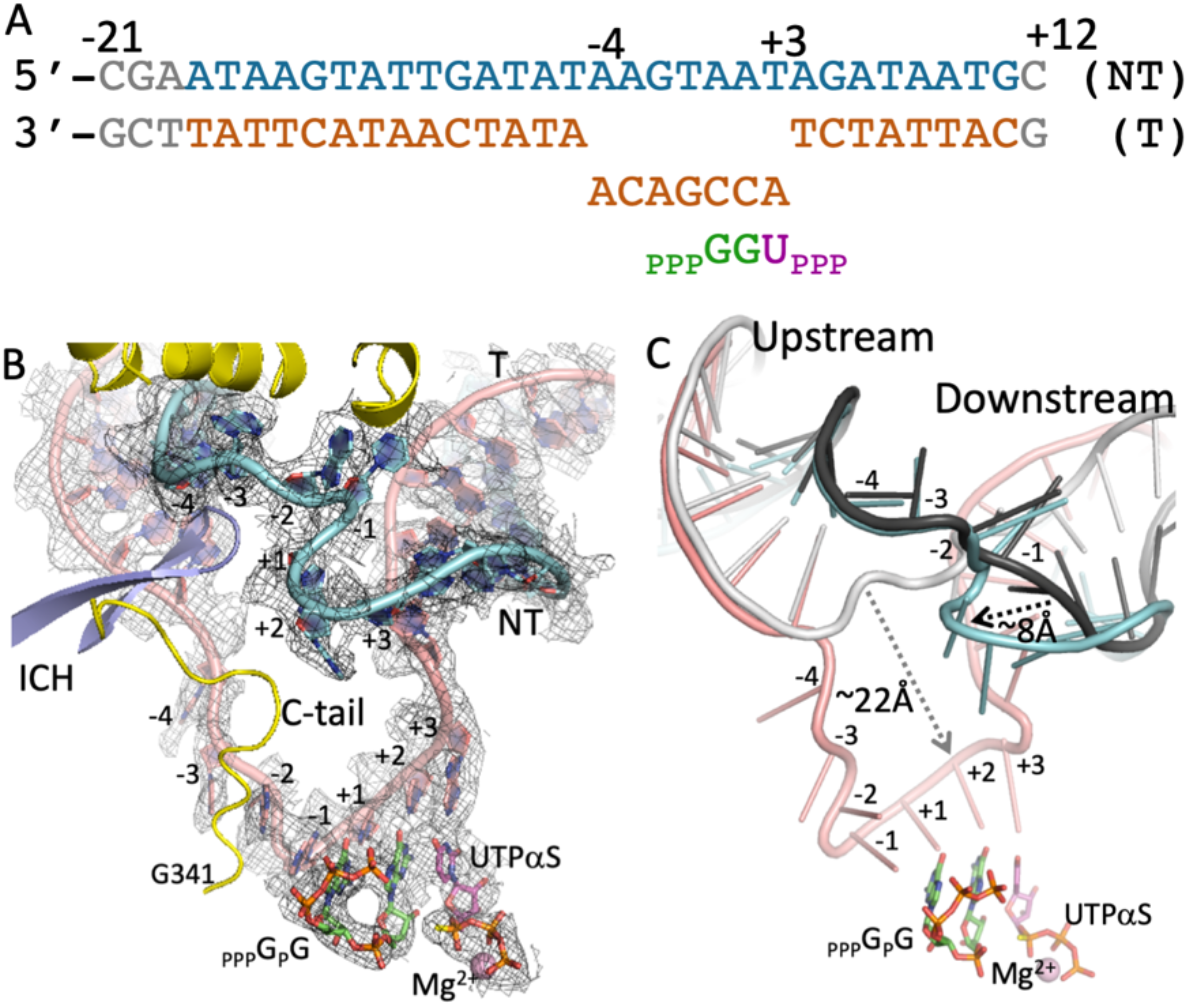
Structure of the yeast mitochondrial RNA polymerase transcription initiation complex (y-mtIC). **A.** The promoter dsDNA template (T, pink) and non-template (NT, cyan), pppGpG RNA (green), and a non-hydrolysable UTPαS (magenta) used in assembling y-mtIC; binding of UTPαS unwinds the downstream +3 base-pair and engages +3 template nucleotide base-paired with UTPαS to form a catalytic-competent quinary IC. **B.** The 3.7 Å resolution density map defines the position and conformation of the transcription bubble, 2-mer RNA, and UTPαS; stereo view, Fig S4. **C.** Superposition of y-mtPIC and y-mtIC shows the large conformational changes (arrows) in the transcription bubble from black NT and gray T strands in y-mtPIC to cyan NT and pink T strands, respectively (Movie S2).

Biochemical studies show that the MTF1 C-tail plays an important role in template alignment, DNA scrunching, and triggering transition into elongation (*15*). The IC structure captures the entire C-tail in the active-site cavity and interacts with the template DNA, NT-loop, and 5’-end of the RNA transcript. The C-tail base is stabilized by ICH, thumb helix, and a loop (521-526) of RPO41 (Fig. 3A). The C-tail tip residue S340 is 4 Å away from the 5’-end α–phosphate of pppGpG RNA. The main chain carbonyls of E338 and H339 hydrogen bond with the N1 and N6 atoms of the −2 template-base; similar interactions with N3 and N4 atoms of unmutated cytosine at −2 position are expected. The C-tail also stabilizes the scrunched NT-loop; M334-Y335 of C-tail stack against the looped-out +1 and +2 NT bases (Fig. 3B). Structural projection indicates that RNA synthesis will progressively push the C-tail out of its position in IC (Fig. 3C), and at a critical length of RNA, the C-tail will be displaced out from the active-site cavity. Single-molecule FRET studies show that IC to EC transition completes at 8-mer RNA synthesis and C-tail deletion delays the transition (*14, 15*). Superposition of y-mtIC on POLRMT EC with 9-bp RNA:DNA (*16*) shows that C-tail must exit for IC to EC transition (Fig. 3D). We expect a similar role of C-tail in homologous h-mtIC (Fig. S6). Upon complete displacement of the C-tail, the MTF1-supporting hairpin will switch its role from guiding the C-tail in IC to supporting the upstream DNA in EC (*16*).

**Figure 3.**
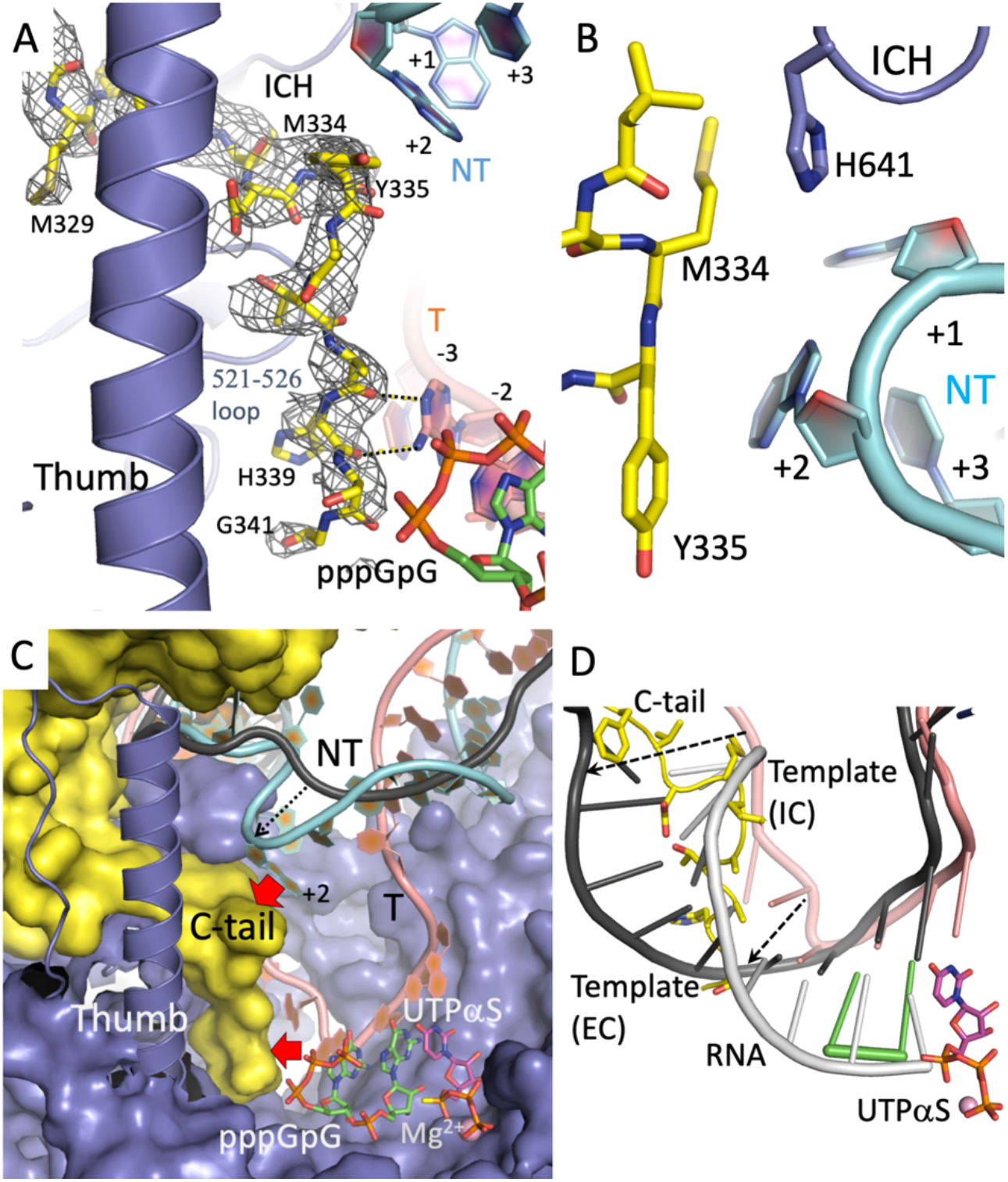
MTF1 C-tail in y-mtIC and its role in stabilizing IC, triggering abortive synthesis, and triggering transition to elongation complex (EC). **A.** The C-tail defined by 3.7 Å map takes a central position in y-mtIC. The C-tail interacts with the intercalating hairpin (ICH), thumb, template (T), non-template (NT), and the RNA transcript. The N1 and N6 of the −2 template nucleotide hydrogen bonds with the main-chain carbonyl groups of E338 and H339, respectively, as shown. **B.** The M334-Y335 platform in C-tail stacks against the +2 NT nucleotide. **C**. The NT strand in y-mtPIC (black) grows towards the C-tail and assumes cyan conformation in y-mtIC. Looping of the scrunched NT strand and the growing RNA:DNA hybrid would push (red arrows) the C-tail to trigger large conformational change for transition from IC to EC. **D.** Superposition of y-mtIC on h-POLMRT elongation complex (EC, PDB Id. 4BOC) shows that transition from IC (pink template and green RNA) to EC (black template and gray RNA) requires ejection of the C-tail.

The PIC and IC structures provide a basis for understanding the mechanism of abortive synthesis. Abortive synthesis is observed in all DNA-dependent RNAPs during transcription initiation, and y-mtRNAP generates large amounts of 2-mer and 3-mer abortive products compared to 4- to 6-mer abortives (Fig. S3F). Two mutually non-exclusive models have been proposed for abortive synthesis: (i) Abortive RNAs are generated by steric clashes between progressively growing RNA:DNA duplex and structural elements such as the N-terminal domain of T7 RNAP (*11*), the C-tail of mitochondrial transcription factors (*15*), or σ3.2/B-finger of multi-subunit transcription factors (*17, 18*). (ii) Abortive RNAs are produced because of scrunching-unscrunching transitions during initial transcription (*19*). Our structures support both mechanisms. The coexistence of scrunched IC and unscrunched PIC is a direct evidence of continuous loading and abortive release of 2-mer RNA in-solution. Steric clashes are predicted between the C-tail and longer RNA:DNA/NT-loop; nascent transcripts will dissociate as abortive products if the C-tail resists to move away. Biochemical studies show that C-tail deletion reduces abortive synthesis (*15*).

The y-mtIC has captured the incoming NTP in a catalytic-competent state poised for incorporation (Fig. 4A), The NTP and the DNA:RNA duplex make extensive interactions with RPO41 in the active site, which is highly conserved in mitochondrial RNAPs (Fig. S7 and S8). The structure of y-mtIC permits reliable modeling of antiviral nucleosides/nucleotides for cytotoxicity prediction. Nucleos(t)ide analogs are widely used to treat viral infections and can cause cytotoxicity by binding to cellular RNAP and mitochondrial POLRMT. Remdesivir is a nucleotide analog with broad antiviral profile (*20*) including treatment of SARS-CoV-2 (COVID-19) infection. Modeling of remdesivir-diphosphate into the NTP-binding pocket of mtRNAP reveals that the characteristic 1’-cyano group of remdesivir clashes with the conserved H1125 in POLRMT (Fig. 4B). Thus, remdesivir is expected to have low cytotoxicity, consistent with its low incorporation efficiency by POLRMT (*20, 21*). Thus, the platform provides a framework for testing mitochondrial toxicity of nucleoside analogs.

**Figure 4.**
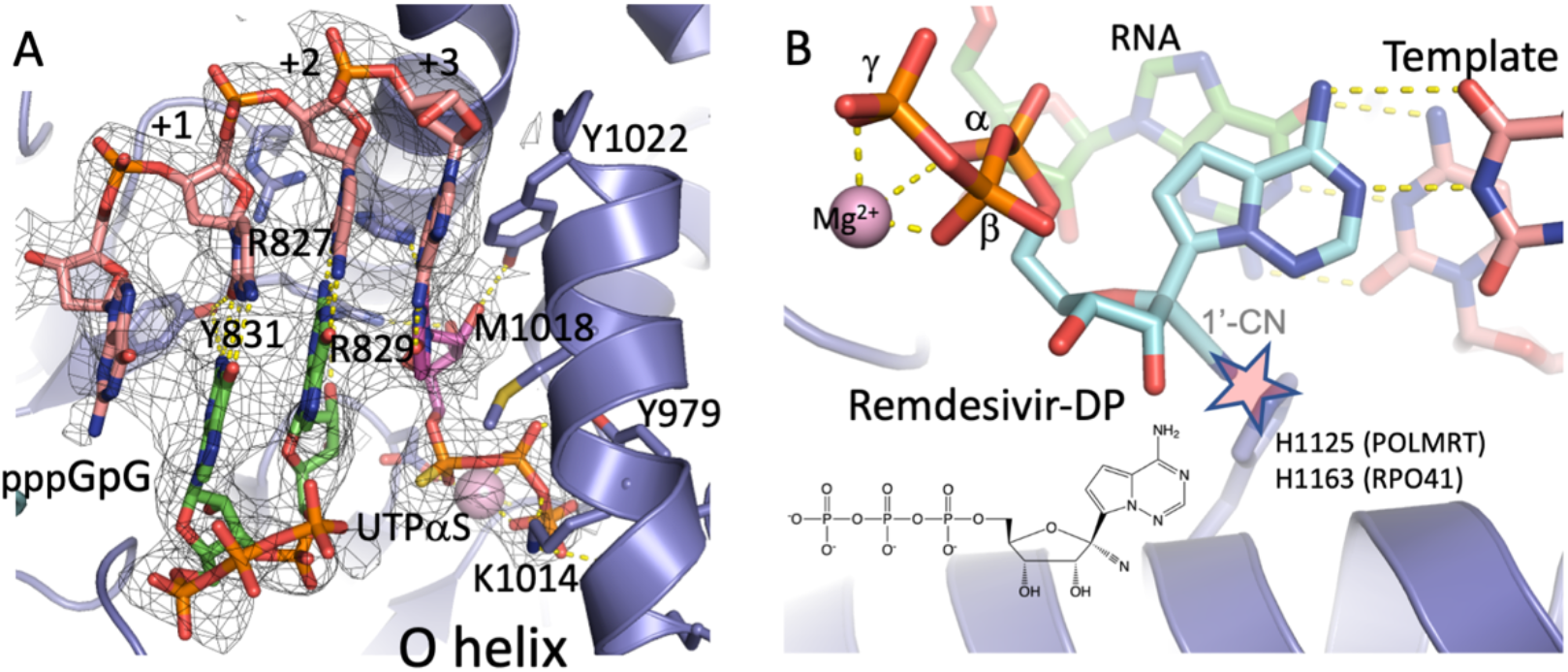
Bound UTPαS and modeled remdesivir-DP in the NTP-binding pocket of y-mtIC. **A.** The 3.7 Å density map of y-mtIC defines the binding of UTPαS in a catalytic competent mode. Key RPO41 residues (POLRMT equivalents in parenthesis) that interact with NTP and RNA:DNA hybrid are listed; R827 (R803), R829 (R805), Y831 (Y807), Y979 (Y956), K1014 (K991), M1018 (M995), and Y1022(Y999). **B.** Modeling shows incompatibility of remdesivir-TP (cyan) binding in the pocket primarily due to a steric clash of the 1’-cyano moiety of inhibitor with the conserved H1125 situated on helix-44 of POLRMT.

## Acknowledgments

We thank Electron Cryogenic Microscopy, Brussels, Thermo Fisher Scientific, Eindhoven, and ESRF-Grenoble for microscope access, and **Marcus Fislage for discussion**.

## Author contributions

B.D.W.; S.S.; H.V.; J.S.; D.V.; S.E.M. investigation and validation; B.D.W; S.S.; S.S.P.; K.D. conceptualization; C.D.; E.K. resources; B.D.W.; E.K.; S.S.P.; K.D. data curation; S.S.P.; K.D. supervision, writing, editing, visualization, funding acquisition, and project administration.

## Data and materials availability

No restrictions on materials, such as materials transfer agreements. The coordinates and density maps for y-mtPIC and y-mtIC structures were deposited under PDB accession numbers 6YMV and 6YMW and EMBD Ids. EMD-10845 and EMD-10846, respectively. All data is available in the main text or the supplementary materials. All data, code, and materials used in the analysis will be available for purposes of reproducing or extending the analysis.

## Supplementary Materials

**Fig. S1:**
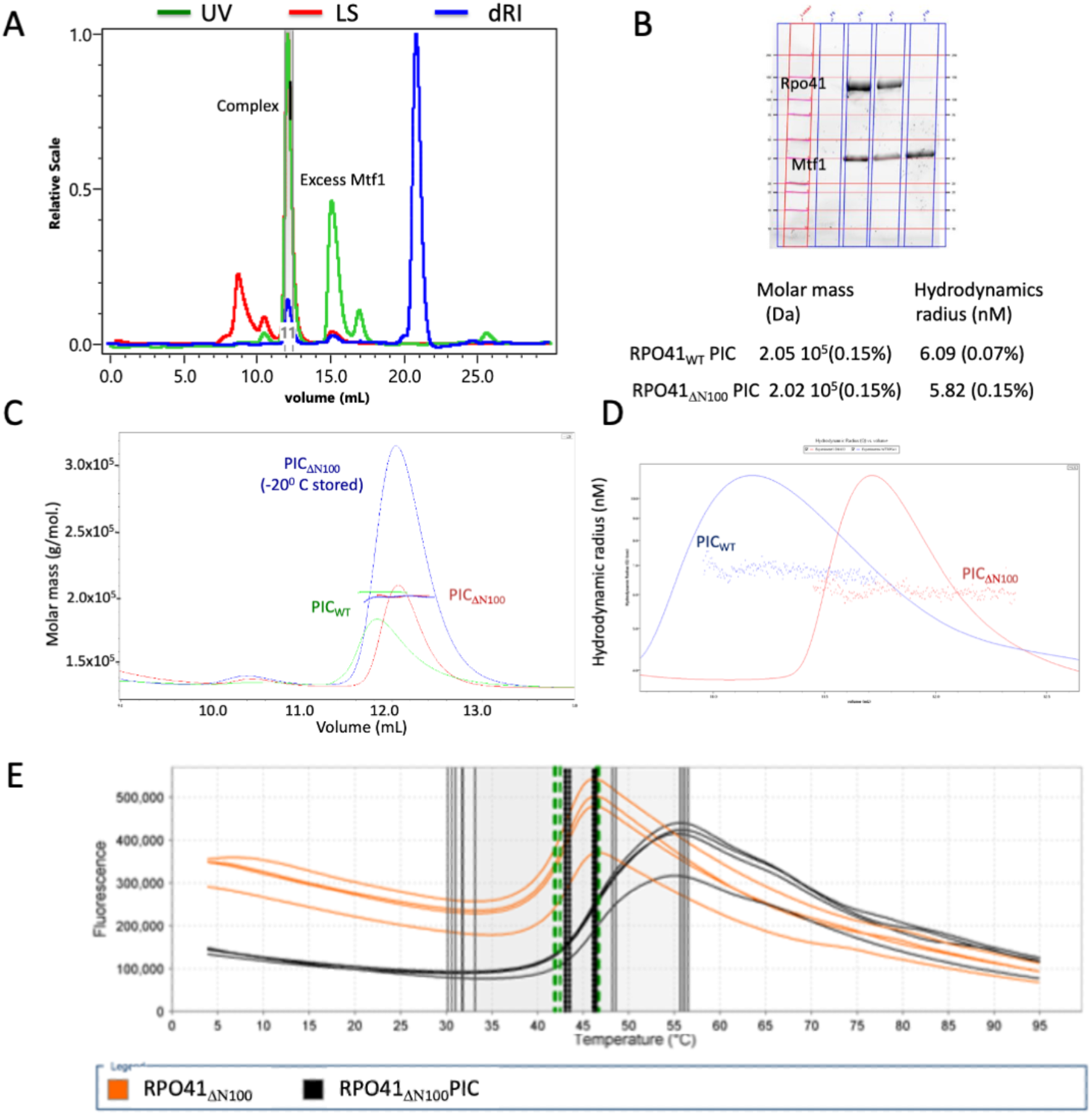
Biophysical characterization of y-mtPIC. (**A**) UV, MALS/DLS, and dRI profile of the y-mtPIC on gel-filtration. (**B**) SDS-PAGE gel of the y-mtPIC fraction (first two from left) and excess MTF1 (right); Calculated molar mass in Dalton (Da) and hydrodynamic radius (nanometer) of y-mtPIC measured by the MALS and DLS setups. (**C**) Comparison of molar mass of y-mtPIC of full-length RPO41 (y-mtPICWT) and of N-terminal 100 amino-acid deletion mutant RPO41 (y-mtPIC_ΔN100_). −80°C freeze-thaw did not damage y-mtPIC_ΔN100_ and the sample could be stored in small aliquots for subsequent uses. (**D**) Hydrodynamic radius measurements of y-mtPICWT vs. y-mtPIC_ΔN100_. (**E**) Protein thermal shift assay profile of y-mtPIC (black curve) vs. RPO41_ΔN100_ (orange curve) in quadruplicate measurements show higher stability of the complex.

**Fig. S2.**
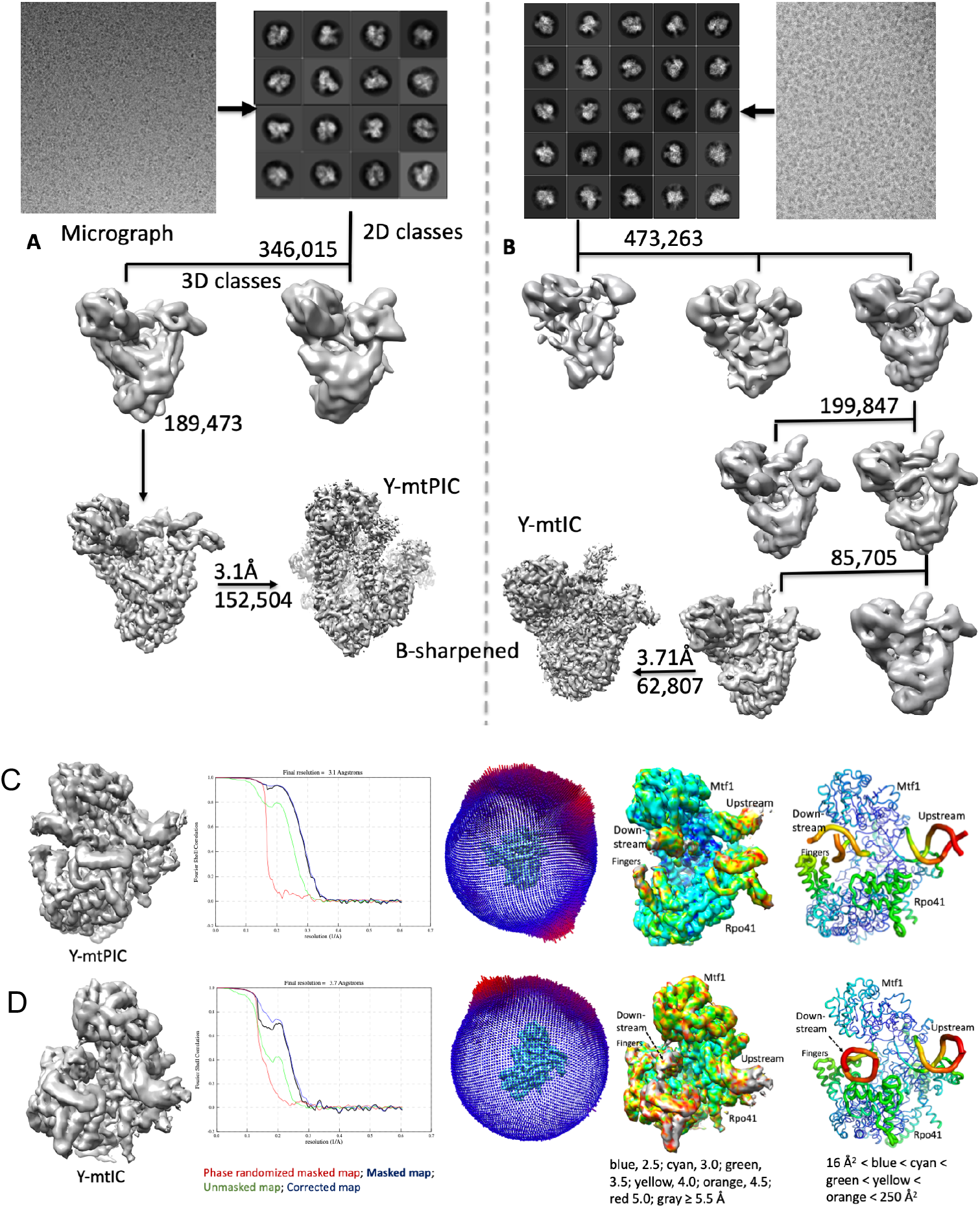
Cryo-EM data processing and density maps. (**A**) Processing of y-mtPIC micrographs and the roadmap for obtaining the final map was calculated at 3.1 Å resolution. The number of particles used at each step are listed. (**B**) Processing of y-mtIC micrographs and the roadmap for obtaining the final y-mtIC map was calculated at 3.7 Å resolution. The y-mtPIC particles grouped in the step 2 of 3D classification were processed to generate a density map at 3.5 Å resolution (not shown here) and map was confirmed to represent a state which is the same as the 3.1 Å y-mtPIC map. For y-mtPIC (**C**) and y-mtIC (**D**) structures, final density map from 3D refinement, FSC curve, angular distribution of the final set of particles, B-sharpened map color coded with estimated local resolution, and the B-factor puffy representation of the final model (blue <cyan<green< yellow< orange colors represent low to high B-factors).

**Fig. S3.**
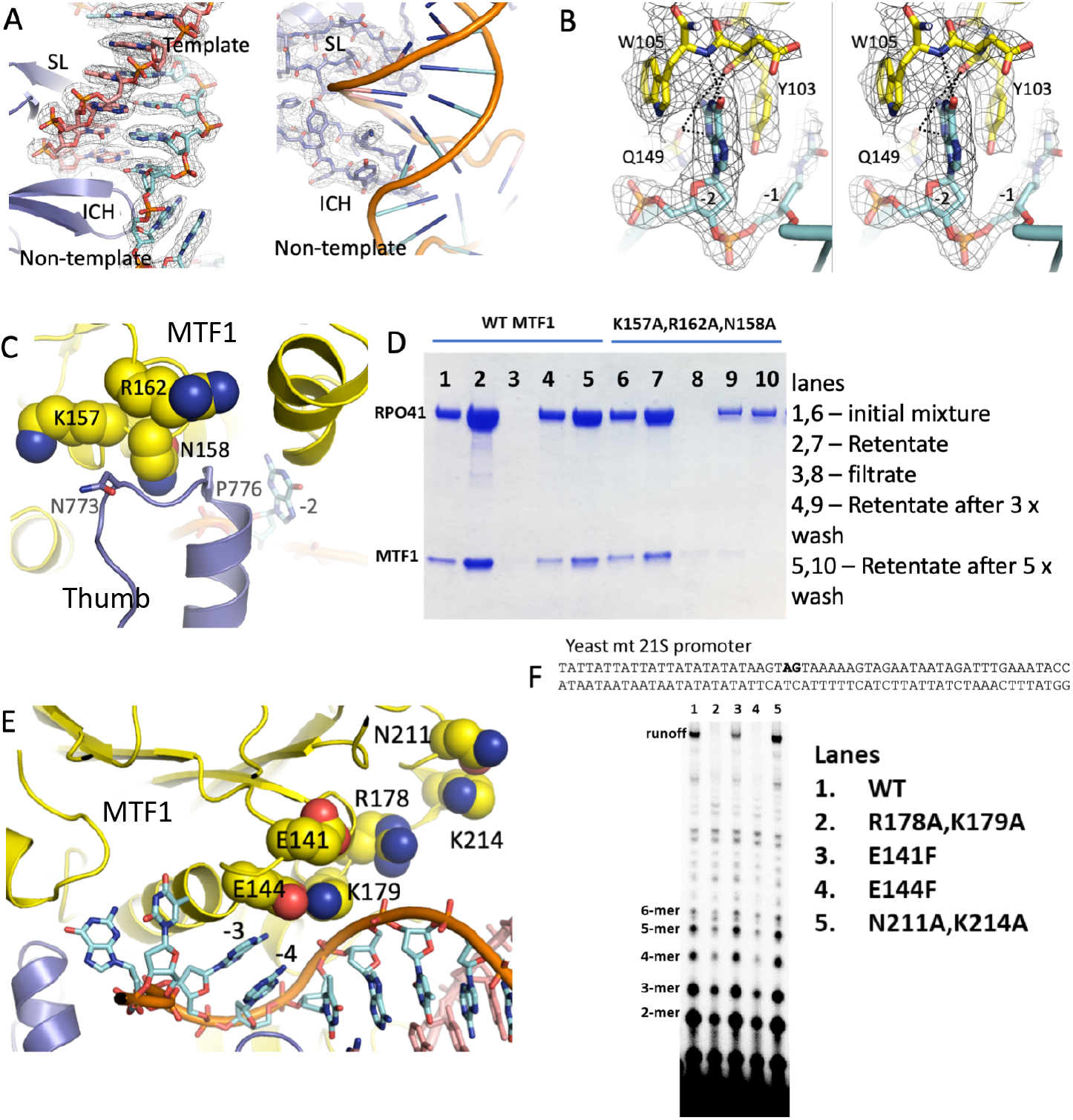
Structural and biochemical analysis of interactions among RPO41, MTF1, and DNA. (**A**) Density for DNA (left) and for intercalating hairpin (ICH) and specificity loop (SL) interacting with DNA (right). (**B**) Stereo view showing the aromatic ring stacking and a hydrogen bond network of −2 NT base with MTF1. (**C**) MTF1 residues K157, N158, and R162 that interact with thumb were mutated to alanine. (**D**) The triple alanine mutant MTF1 was investigated for complex formation with RPO41 by an ultrafiltration assay. Lane 5 shows a stable RPO41-MTF1 complex after five washes whereas the mutant MTF1 is washed out as shown in lane 10. (**E**) MTF1 sites in and around NT grove that were mutated to ascertain their role in polymerization activity. (**F**) Sequence of the yeast mitochondrial *21S* rRNA promoter (−25 to +32) with start-site in bold. Image of the polyacrylamide gel (24% polyacrylamide 4 M Urea denaturing gel) show the abortive and runoff RNA products of the transcription reactions carried out with 1 μM Rpo41, 2 μM Mtf1 and 2 μM promoter duplex for 15 min using 100 μM ATP, UTP, GTP, and 1.25 mM 3’-dCTP spiked with γ[^32^P]ATP. K179 interacts with the −5 position promoter NT, and the double mutant (R178A + K179A) MTF1 abrogates runoff synthesis. E144 interacts with the −3 NT nucleotide and its mutation to phenylalanine abrogates RNA synthesis. Mutations of the remaining amino acids that are not near the DNA have little effect on runoff synthesis.

**Fig. S4.**
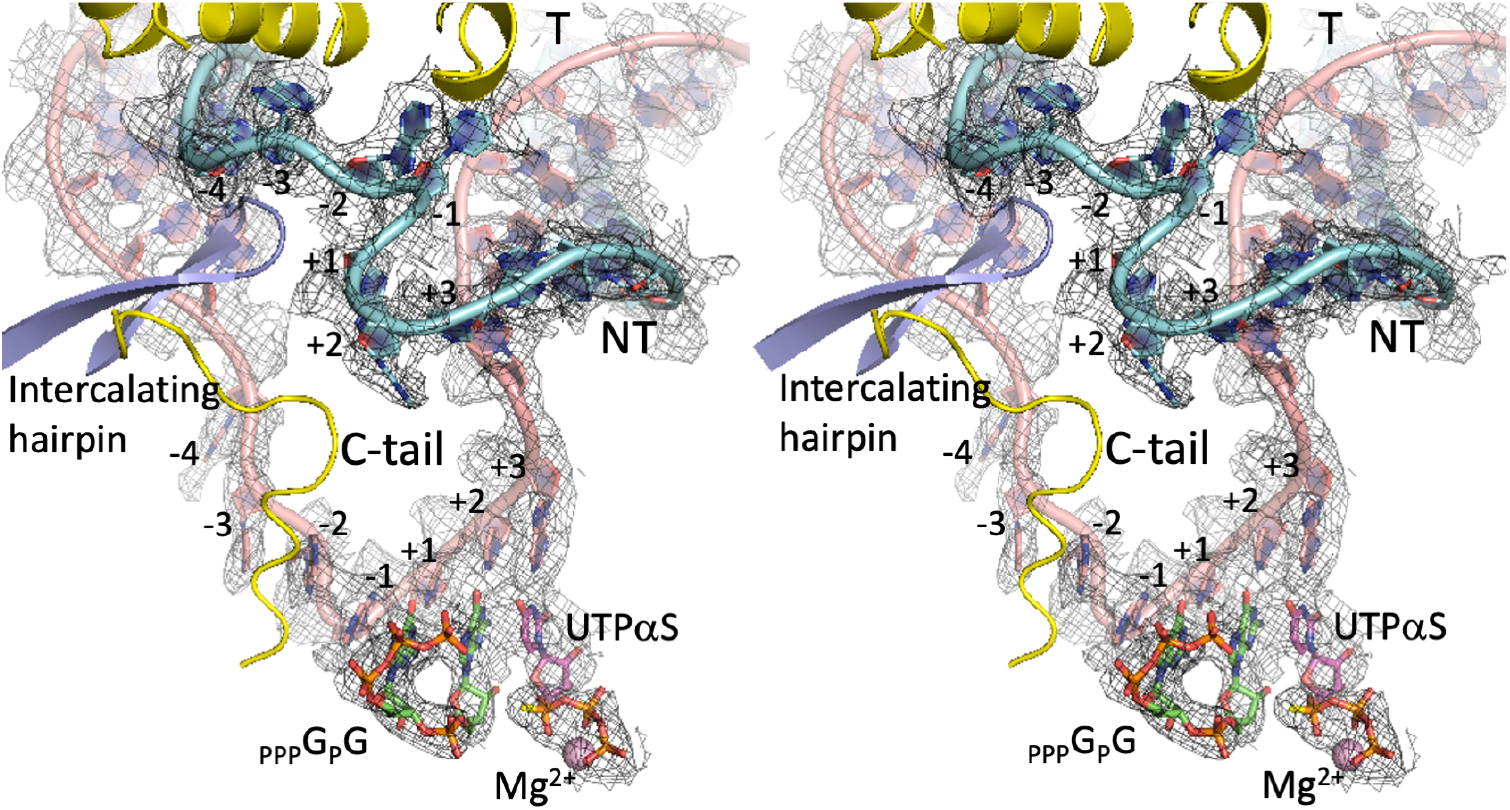
Stereo view of the transcription bubble in y-mtIC. The cryo-EM density map at 3.7 Å resolution help build the DNA reliably. Most protein atoms are omitted for a clear view. The non-template (NT; cyan) and template (T; pink) nucleotides in the bubble are numbered. The pppGpG (RNA) and UTPαS are in green and magenta C-atom representations, respectively.

**Fig. S5.**
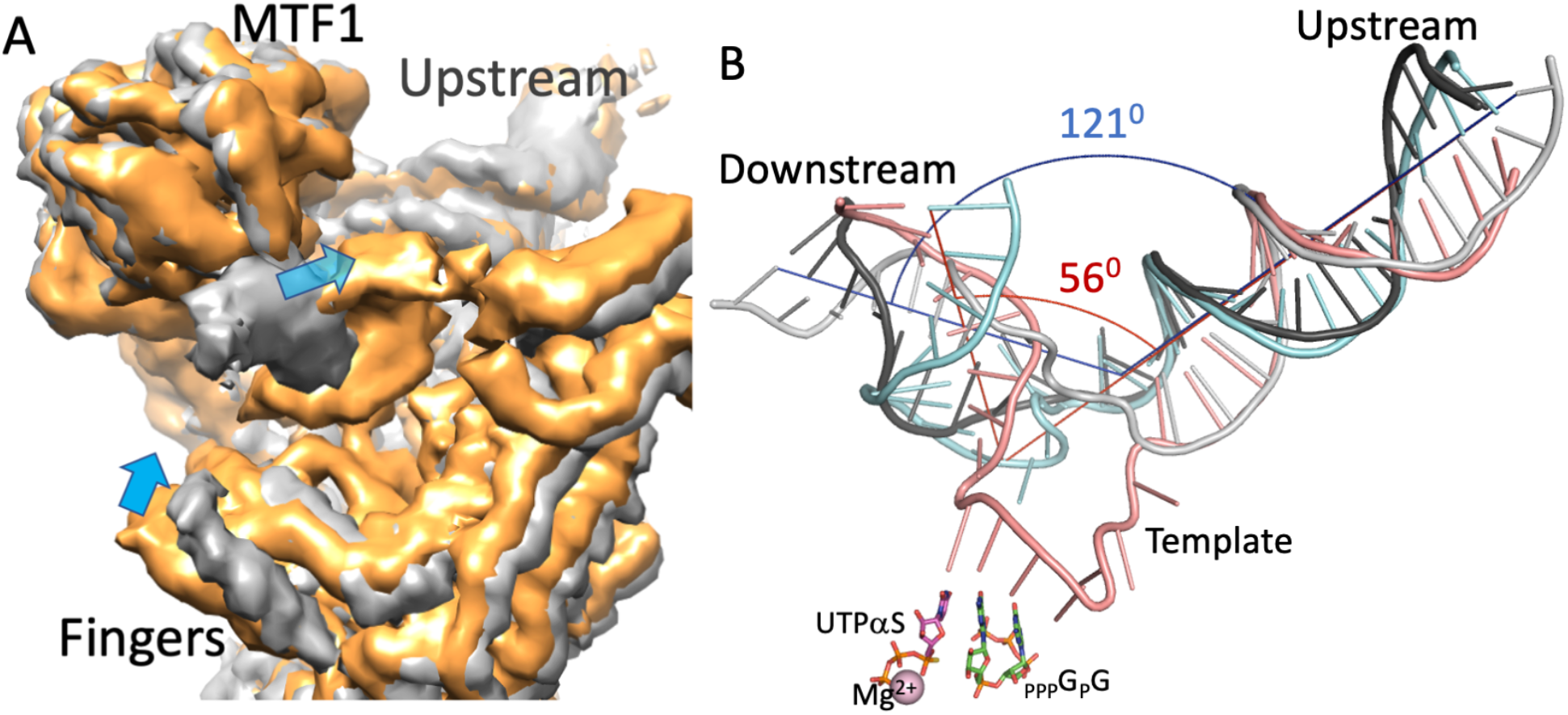
Bending of downstream DNA during PIC to IC transition. (**A**) Comparison of density maps of y-mtPIC (gray) and y-mtIC (orange) show that the downstream DNA bends further and fingers close in as PIC switches to IC; the blue arrows indicate the motion of downstream DNA and fingers. (**B**) Superposition of y-mtPIC structure (black non-template; gray template) on y-mtIC structure (cyan non-template; pink template) shows the bending of downstream DNA while the upstream DNA in both structures are aligned. The angle between the upstream and downstream DNA are about 120° and 60°, respectively, in y-mtPIC (blue axes) and y-mtIC (red axes); i.e., the DNA is bent by ~60° in PIC and subsequently by another 60° to ~120° in the IC structure. The DNA bending calculations were done using CURVES+ server (*11*). Movies S1-S3 shows the conformational changes during the transition from PIC to IC state.

**Fig. S6.**
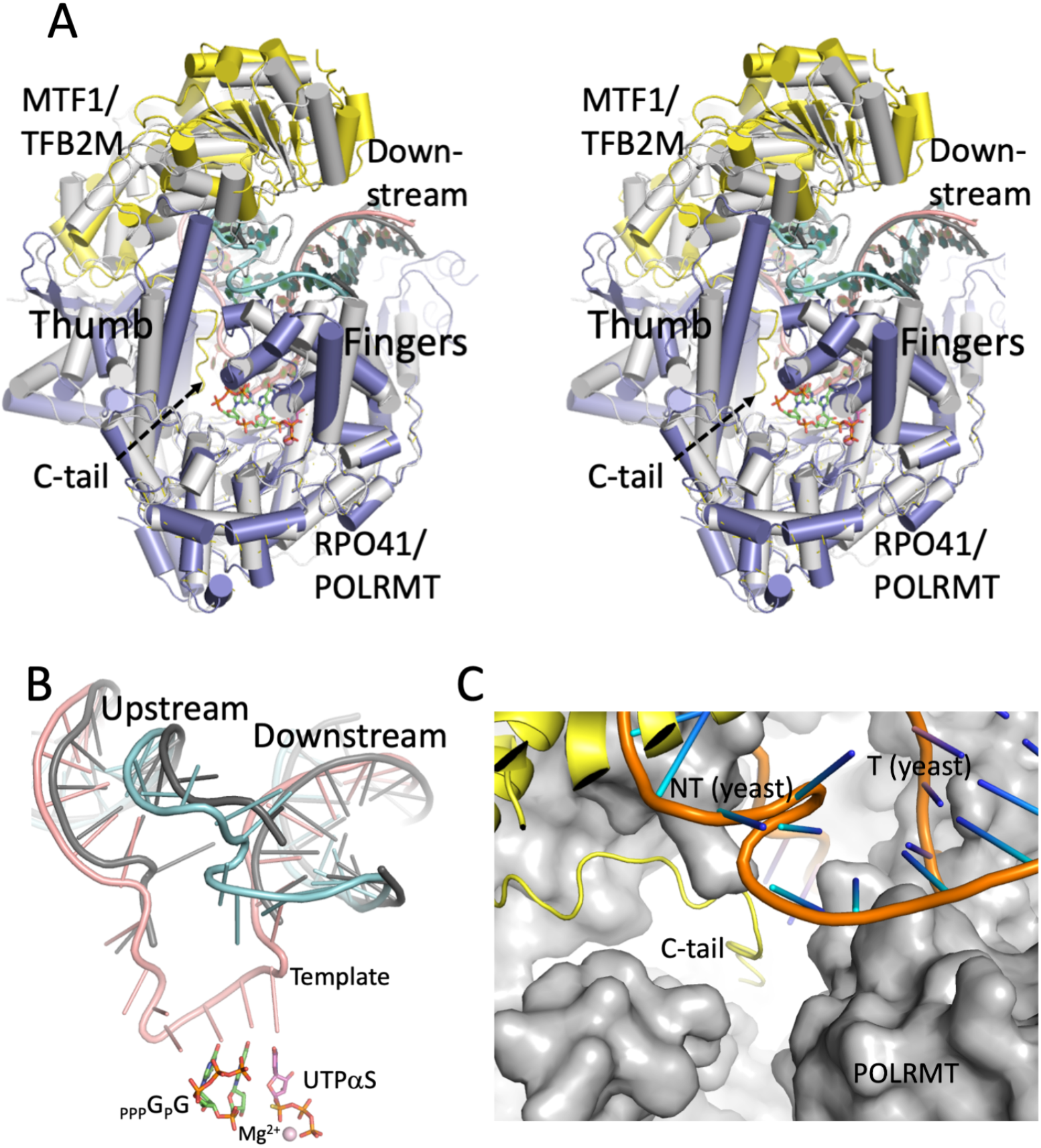
Comparison of y-mtIC and h-mtIC structures. (**A**) A stereo view of the overlaid y-mtIC structure on h-mtIC (PDB ID. 6EQR) based on superposition of RPO41 and POLRMT. The h-mtIC components POLMRT and TFB2M are in gray and DNA in dark gray. The y-mtIC components RPO41, MTF1, non-template and template are in blue, yellow, cyan, and pink, respectively. (**B**) The tracks of the upstream and downstream DNA in both structures are well superimposed. (**C**) Based on the above superposition, the C-tail of y-mtIC is positioned in the active-site cavity of h-mtIC suggesting that the TFB2M C-tail has an analogous role in h-mtIC to the role of C-tail in y-mtIC.

**Fig. S7.**
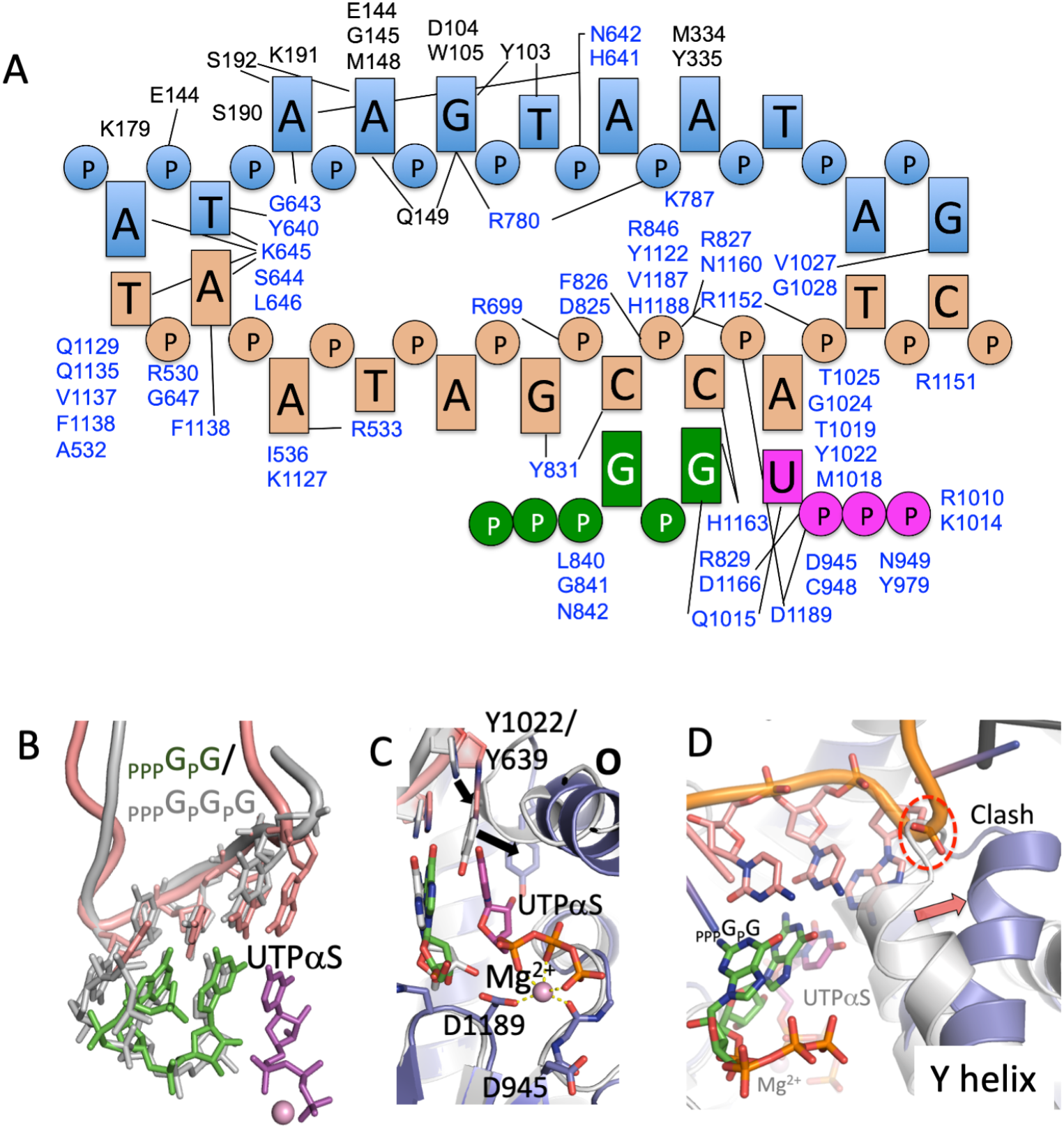
Interactions of transcription bubble in y-mtIC structure and comparison of active site region with T7 RNAP IC and h-mtRNAP IC. (**A**) A schematic representation of the interactions of the transcription bubble with surrounding protein residues of MTF1 (black) and RPO41 (blue); the non-template strand, template strand, pppGpG (RNA) and UTPαS are in cyan, light brown, green, and magenta, respectively. (**B**) A comparison of y-mtIC with T7 RNAP IC structures (PDB ID. 1QLN) shows that the promoter DNA template in y-mtIC (pink) is bent sharply at the active site and after 4 nucleotides that is analogous to the template track in T7 RNAP IC (gray); however, the bound UTPαS captures y-mtIC in the catalytic mode for nucleotide incorporation whereas, the T7 RNAP IC structure represents the post-translocated state with no bound NTP. (**C**) The comparison also shows that the NTP-binding pocket undergoes a conformational change. The conserved Y639 on the O helix of T7 RNAP must shift to the position that Y1022 of RPO41 takes to accommodate an NTP. (**D**) Superposition of y-mtIC on h-mtIC structure (PDB Id. 6EQR) shows that the Y-helix of POLRMT clashes with the template strand and the state observed in the h-mtIC structure would not position the template in a conformation that is compatible for RNA/NTP-binding. The h-mtIC structure represents an inactive-clenched state. The Y-helix (1023-1041 of RPO41) is critical for downstream DNA unwinding.

**Fig. S8.**
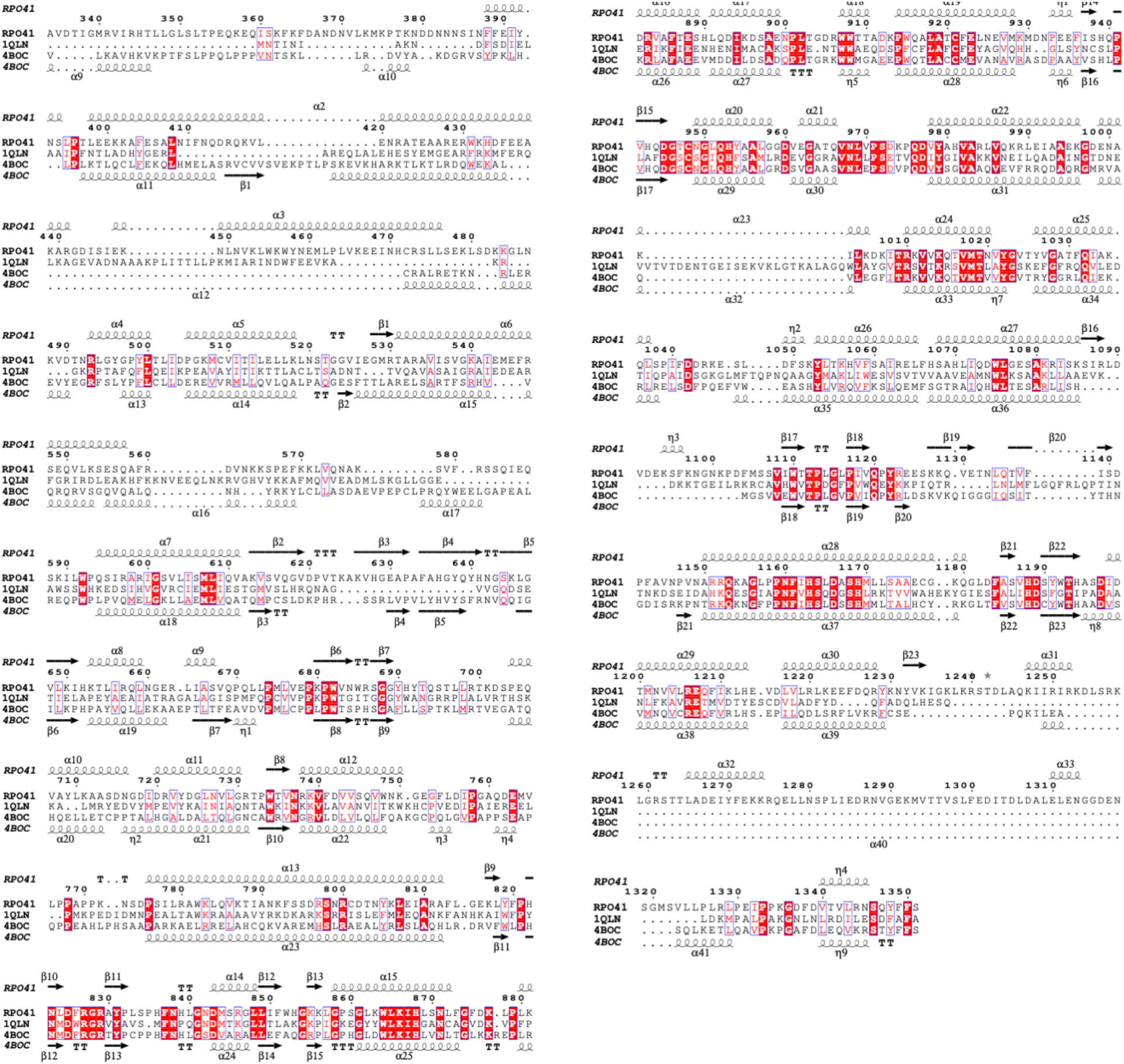
Amino acid sequence alignment of RPO41, T7 RNAP (PDB Id. 1QLN), and human POLRMT (PDB Id. 4BOC). The secondary structures of RPO41 and POLRMT are shown on top and bottom of the aligned sequences, respectively, using the server ESPript 3.0 (http://espript.ibcp.fr/ESPript/cgi-bin/ESPript.cgi).

**Table S1.** Cryo-EM data and structure analysis statistics.

|  |  |  |
| --- | --- | --- |
| Structure | y-mtPIC (RPO41/ MTF1/dsDNA) | y-mtIC (RPO41/MTF1/<br>dsDNA/RNA/UTPαS) |
| PDB Id/EMBD Id | 6YMV/EMD-10845 | 6YMW/EMD-10846 |
| Data collection |  |  |
| Grid type | Quantifoil R1.2/1.3 | Quantifoil R1.2/1.3 |
| Number of grids | 1 | 1 |
| Microscope/detector | Titan Krios/Gatan K2 | Titan Krios/Gatan K2 |
| Voltage (kV) | 300 | 300 |
| Magnification | 165,000 x | 165,000 x |
| Recording mode | Counting | Counting |
| Dose (e<sup>-</sup>/Å<sup>2</sup>/frame) | 1.09 | 1.22 |
| Total dose (e/Å<sup>2</sup>) | 65 | 61 |
| Number of frames/movie | 60 | 50 |
| Total exposure time (sec) | 6 | 5 |
| Pixel size (Å) | 0.827 | 0.827 |
| Defocus range (Å) | -7000 to - 22000 | -7000 to - 22000 |
| Data processing |  |  |
| Number of micrographs used | 1,302 | 2,684 |
| Number of particles picked | 492,574 | 775,698 |
| Number of particles after 2D | 346,015 | 473,263 |
| Particles used for final map | 152,504 | 62,807 |
| Map resolution (FSC 0.143; Å) | 3.1 | 3.7 |
| Map sharpening B factor (Å<sup>2</sup>) | -114.98 | -163.95 |
| Model fitting |  |  |
| Experimental map/model correlation | 0.84 | 0.79 |
| Total number of atoms | 10,944 | 11,353 |
| Average B factor (Å<sup>2</sup>) |  |  |
| Clash score | 11.2 | 13.1 |
| Ramachandran plot; favored/outlier (%) | 95.79/0.08 | 94.69/0.00 |
| Rotamer outlier (%) | 0.0 | 0.27 |
| RMSD bond length (Å<sup>2</sup>)/ bond angle (°) | 0.007/0.728 | 0.008/0.81 |

## Movies S1 - S3

**Movie S1. Overall structural change in the transition from PIC to IC.** Morphing between the transcription pre-initiation state (y-mtPIC, gray RPO41 and DNA, yellow MTF1) and initiation state (y-mtIC, yellow MTF1, blue RPO41, cyan non-template, and pink template) simulates the conformational changes in the promoter and protein during transition from the PIC to IC state. The downstream DNA bends inward and parts of the C-terminal domain including fingers (in front) undergo large conformational changes. MTF1, upstream DNA, and parts of the N-terminal domain of RPO41 that interact with MTF1 and upstream DNA show minimal conformational changes; e.g. the thumb helix on the left and MTF1 on the top have minimal movements.

**Movie S2. Conformational change of downstream DNA and expansion of the transcription bubble.** Morphing between the PIC state (gray DNA) and IC state (cyan non-template, pink template, and 2-mer RNA pppGpG and UTPαS in stick models) shows DNA bubble expansion associated with the template base-pairing with the 2-mer RNA and UTP at the polymerase active site. The downstream DNA bends by about 60**°**(Fig. S5). The protein atoms are removed for clear visualization of the DNA.

**Movie S3. Scrunching of the non-template DNA strand as an NT-loop.** Morphing between the PIC state (gray DNA) and IC state (cyan non-template, pink template, and 2-mer RNA pppGpG and UTPαS in stick models) shows looping of the non-template strand into an NT-loop. This looping appears to be a major contributor to bending of the downstream DNA with respect to the upstream DNA (Fig. S5).

